# SpLitteR: Diploid genome assembly using TELL-Seq linked-reads and assembly graphs

**DOI:** 10.1101/2022.12.08.519233

**Authors:** Ivan Tolstoganov, Zhoutao Chen, Pavel A. Pevzner, Anton Korobeynikov

## Abstract

**Background:** Recent advances in long-read sequencing technologies enabled accurate and contiguous *de novo* assemblies of large genomes and metagenomes. However, even long and accurate high-fidelity (HiFi) reads do not resolve repeats that are longer than the read lengths. This limitation negatively affects the contiguity of diploid genome assemblies since two haplomes share many long identical regions. To generate the telomere-to-telomere assemblies of diploid genomes, biologists now construct their HiFi-based phased assemblies and use additional experimental technologies to transform them into more contiguous diploid assemblies. The barcoded linked-reads, generated using an inexpensive TELL-Seq technology, provide an attractive way to bridge unresolved repeats in phased assemblies of diploid genomes.

**Results:** We developed SpLitteR tool for diploid genome assembly using linked-reads and assembly graphs and benchmarked it against state-of-the-art linked-read scaffolders ARKS and SLR-superscaffolder using human HG002 genome and sheep gut microbiome datasets. The benchmark showed that SpLitteR scaffolding results in 1.5-fold increase in NGA50 compared to baseline LJA assembly and other scaffolders while introducing no additional misassemblies on the human dataset.

**Conclusion:** We developed the SpLitteR tool for haplotype phasing and scaffolding in an assembly graph using barcoded linked-reads. We benchmarked SpLitteR on assembly graphs produced by various long-read assemblers and have shown how TELL-Seq reads facilitate phasing and scaffolding in these graphs. This benchmarking demonstrates that SpLitteR improves upon the state-of-the-art linked-read scaffolders in the accuracy and contiguity metrics. SpLitteR is implemented in C++ as a part of the freely available SPAdes package and is available at https://github.com/ablab/spades/releases/tag/splitter-preprint.

## Background

The recently developed linked-read technologies, such as stLFR [1], TELL-Seq [2], and LoopSeq [3], are based on co-barcoding of short reads from the same long DNA fragment. They start with the distribution of long DNA fragments over a set of containers marked by a unique barcode. Afterward, long fragments within the containers are sheared into shorter fragments and sequenced. The resulting library consists of linked-reads, and short reads marked by the barcode corresponding to the set of long fragments.

Various tools, such as Athena [4], cloudSPAdes [5], Supernova [6], and TuringAssembler [2], were developed to generate *de novo* genome assembly from linked-reads alone. However, even though linked-reads result in more contiguous assemblies than assemblies based on non-linked short reads, all these tools generate rather fragmented assemblies of large genomes and metagenomes. For large genomes and metagenomes, long high-fidelity (HiFi) reads proved to be useful in generating highly-accurate and contiguous assemblies [7–12]. Still, even though HiFi reads enabled the first complete assembly of the human genome by the

Telomere-to-Telomere (T2T) consortium [13], HiFi assemblies do not resolve some long repeats and thus are often scaffolded using supplementary technologies, such as Hi-C reads, Oxford Nanopore (ONT) ultralong reads, and Strand-seq reads [13]. Scaffolding methods based on inexpensive linked-reads represent a viable alternative to other supplementary technologies since they combine the low cost of short reads and the long-range information encoded by linked-reads originating from the same barcoded fragment.

Although the state-of-the-art linked-read scaffolders, such as Architect [14], ARKS [15], Physlr [16], and SLR-superscaffolder [17] improve the contiguity of HiFi assemblies, they do not take advantage of the assembly graph and thus ignore the important connectivity information encoded by this graph. In addition, these tools are not applicable to diploid assemblies and complex metagenomes with many similar strains.

We present the SpLitteR tool that uses linked-reads to improve the contiguity of phased HiFi assemblies. In contrast to existing linked-reads scaffolders, it utilizes the assembly graph and was developed with diploid assemblies in mind. Given a linked-read library and a HiFi assembly graph in the GFA format, SpLitteR resolves repeats in the assembly graph using linked-reads and generates a simplified (more contiguous) assembly graph with corresponding scaffolds.

## Results

We benchmarked SpLitteR on three different datasets.

HUMAN dataset [2] was obtained from a diploid human HG002 genome that was recently assembled from HiFi reads [12]. The HUMAN dataset includes a TELL-Seq library which contains ∼994 million barcoded TELL-Seq reads and a HiFi read-set from HG002. Since both TELL-Seq [2] and HiFi technologies [18] emerged only three years ago, there are currently very few datasets that include both HiFi and TELL-Seq reads. We thus generated additional TELL-Seq datasets described below.

HUMAN+ dataset includes two additional TELL-Seq libraries which contain an additional ∼4,585 million barcoded TELL-Seq reads.

The SHEEP dataset includes a TELL-Seq library containing ∼1004 million barcoded reads and a HiFi library from a sheep fecal metagenome. Table 1 provides information about these datasets, such as approximate fragment length. The Data Preparation section specifies the details of the TELL-Seq library preparation.

**Table 1.** Information about TELL-Seq datasets. Mean fragment length was evaluated based on the T2T reference for the HUMAN dataset, and the metaFlye [9] assembly for the SHEEP dataset. A fragment is defined as the set of multiple paired-end reads with the same barcode aligned to the same long (>500kbp) scaffold. Mean fragment length is the mean distance from the start of the leftmost aligned read in a fragment to the end of the rightmost aligned read in a fragment.

| Dataset | Number of TELL-Seq reads | Genome coverage | Mean fragment length |
| --- | --- | --- | --- |
| HUMAN | 993,847,904 | 47x | 34,317 |
| HUMAN+ | 5,579,154,072 | 267x | 36,211 |
| SHEEP | 1,005,083,124 | N/A | 27,719 |

SpLitteR (version 0.1) was benchmarked against ARKS 1.2.4 [15], and SLR-superscaffolder 0.9.1 [17] on the HUMAN, HUMAN+, and SHEEP datasets. We were not able to run Physlr [16] on TELL-Seq data, while Architect [14] was not able to finalize the scaffolding step after 14 days of runtime. We used LJA v0.2 [11] to generate the assembly graph (multiplex de Bruijn graph) from HiFi reads in the HUMAN and HUMAN+ datasets, and metaFlye (v.2.9) [9] to generate the assembly graph for the SHEEP dataset. Assemblies for both datasets were further scaffolded using SpLitteR, ARKS, and SLR-superscaffolder. We used QUAST-LG [19] to compute various metrics of the resulting assemblies (NGA50 values, the largest alignment, etc.) with the homopolymer-compressed T2T HG002 assembly as the reference [12] for the HUMAN and HUMAN+ datasets.

### HUMAN dataset benchmark

We benchmarked SpLitteR, ARKS 1.2.4 [15], and SLR-superscaffolder 0.9.1 [17] on the HUMAN dataset. In the case of SpLitteR, we benchmarked an assembly formed by sequences of homopolymer-compressed edges in the simplified assembly graph generated by the SpLitteR. In the cases of ARKS and SLR-superscaffolder, we benchmarked their assemblies formed by scaffolds of the sequences of the LJA-generated edges.

We used QUAST-LG [19] to compute various metrics of the resulting assemblies with the homopolymer-compressed T2T HG002 assembly as the reference [12]. Table 2 illustrates that SpLitteR resulted in the largest NGA50 and NGA25 metrics for the HUMAN dataset. Specifically, NGA50 values are 301, 303, 301, and 461 kb for LJA (input graph), ARKS, SLR-superscaffolder, and SpLitteR, respectively. For the HUMAN+ dataset, the LJA assembly scaffolded with SpLitteR resulted in a 479 kb NGA50 value. Reduced total length for SpLitteR is explained by glueing together edges adjacent to resolved vertices, as the length of the vertex was included in the total length in the original LJA assembly for both in-edge and out-edge. The vertex length in the LJA graph that was used for benchmarking can reach 40 kbp. Since ARKS is using non-unique k-mers for its barcode-to-contig assignment procedure, pairs of consecutive edges originating from the same haplotype and from different haplotypes have roughly the same number of shared barcodes, which prevents accurate phasing. This might explain roughly the same NGA50 metric and higher number of misassemblies for ARKS scaffolding compared to baseline LJA assembly. SLR-superscaffolder was not able to locate unique contigs in the assembly, and thus produces the assembly identical to LJA.

**Table 2.**
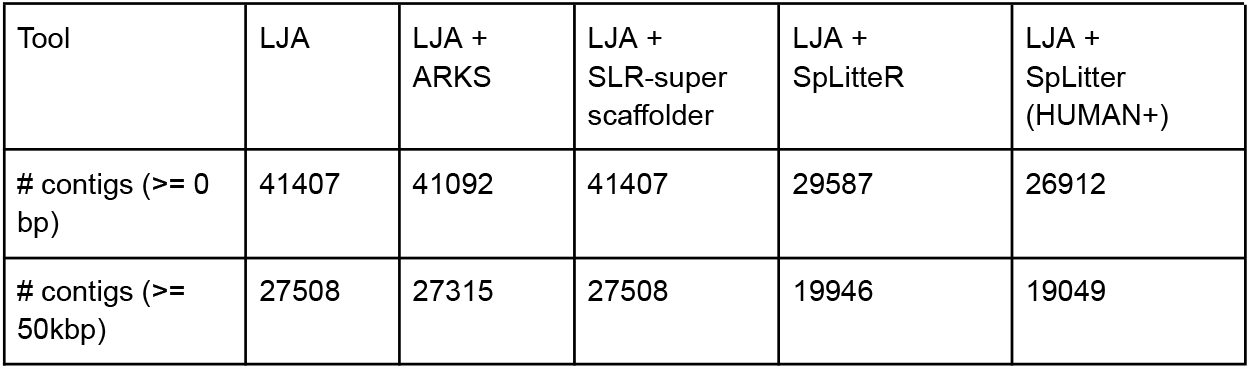
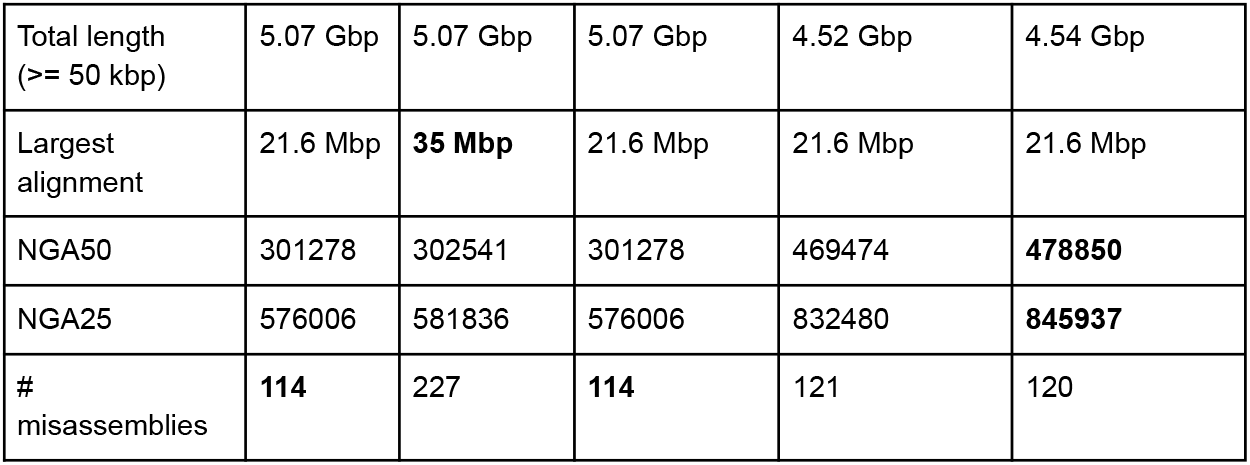
QUAST results for the HUMAN dataset.

ARKS generated the longest misassembly-free scaffold of length 35 Mbp (as compared to 21.6 Mbp for SpLitteR). The largest ARKS scaffold comprises three long edges of the LJA assembly graph aligned to human chromosome X. These edges (Figure 1, blue edges) are divided by two bubbles of length approximately 80 kbp (Figure 1, red edges). Closer investigation showed the central blue edge to be a graph construction artifact, since two halves of the edge were aligned by QUAST-LG to different haplotypes. As a result, the repeat shown at Figure 1 could not be resolved by SpLitteR, as it does not contain any branching vertices incident to the central blue edge (SpLitteR only attempts to resolve repeats corresponding to branching vertices, i.e., vertices with both indegree and outdegree exceeding 1).

**Figure 1.**
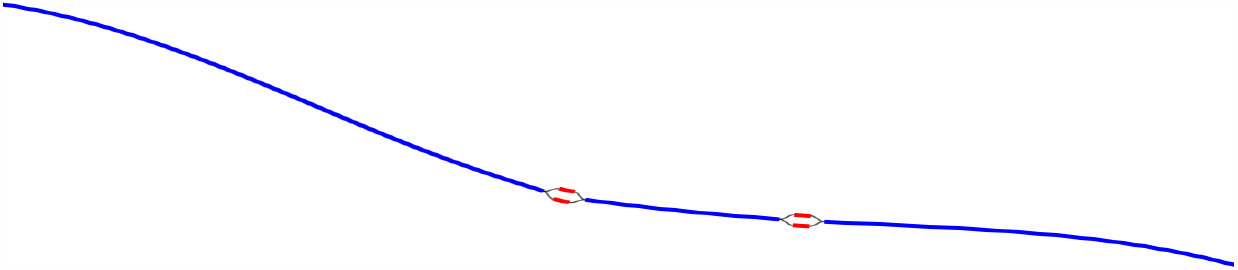
Bandage-NG plot of the ARKS largest scaffold. Three edges comprising the largest ARKS scaffold are shown in blue. Bulge edges are shown in red. The absence of branching vertices makes it impossible for SpLitteR to resolve this component.

#### Repeat classification results

SpLitteR repeat resolution algorithm processes every *repeat* vertex with at least 2 in-edges and 2 out-edges in the assembly graph individually. We classify repeat vertices based on the results of the repeat resolution procedure. Given a repeat vertex *v*, we say that a pair (*in_edge, out_edge)* of in- and out-edges of *v* is *connected*, if *out_edge* follows *in_edge* in the genomic path according to the repeat resolution procedure. Repeat vertex *v* is *partially resolved*, if there is at least one connected pair of edges, *completely resolved* if all in- and out-edges of *v* belong to a connected pair, and *uncovered* if there is no in-edge and out-edge pair of edges that share at least 2 barcodes. If *v* is not completely resolved, partially resolved, or uncovered, we say that *v* is *ambiguous*. In the case of a diploid assembly, completely and partially resolved vertices correspond to regions of the assembly that were phased. For the partially resolved vertices, only one of the haplotypes was recovered using the barcode information, while the other was recovered by the process of elimination. Repeat vertex classification is described in more detail in the “Repeat resolution” section.

Figure 2 shows the number of completely resolved, partially resolved, ambiguous, and uncovered vertices for the HUMAN dataset depending on the length of the repeat vertex. The total number of completely resolved, partially resolved, ambiguous, and uncovered vertices for the HUMAN, HUMAN+, and SHEEP datasets is shown in Table 3. The relatively high number of ambiguous vertices in the SHEEP dataset can be explained by higher mean in- and out-degrees of vertices in the contracted metaFlye assembly graph.

**Table 3.** Repeat resolution statistics for the HUMAN, HUMAN+, and SHEEP datasets.

| Dataset | # Completely resolved vertices | # Partially resolved vertices | # Uncovered vertices | # Ambiguous vertices |
| --- | --- | --- | --- | --- |
| HUMAN | 7202 | 2853 | 6241 | 547 |
| HUMAN+ | 10681 | 2405 | 3526 | 231 |
| SHEEP | 144 | 424 | 517 | 421 |

**Figure 2.**
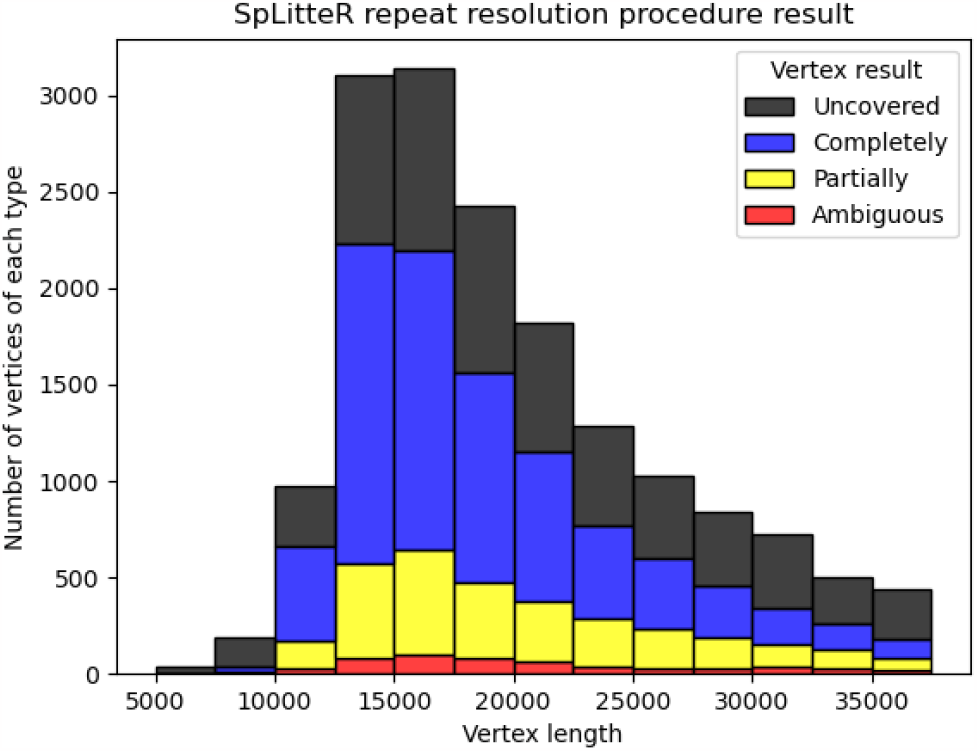
Information about repeat resolution for the HUMAN dataset. The x-axis shows the length of the vertex (approximate length of a repeat) in the LJA assembly graph. Barplots show the number of completely resolved (blue), partially resolved (yellow), uncovered (black), and ambiguous (red) vertices identified by the SpLitteR repeat resolution procedure. The bin width in the histogram is 2500 bp.

#### Trio-binning validation

We additionally used a *trio-binning* tool LJATrio [20], which employs the parental mother/father Illumina short reads to validate the repeat resolution procedure for a diploid dataset from a child. LJATrio uses trio information to classify edges of the multiplex de Bruijn graph (constructed from the HiFi reads from the child dataset) into maternal, paternal, or undefined. We applied LJATrio to the mother-father-child dataset where the child corresponds to the HG002 genome and classified all edges of the corresponding contracted assembly graph into maternal, paternal, or undefined. For each vertex *v*, we classify its resolved in- and out-edge links as *correct* if both edges of the link are marked as either paternal or maternal, *incorrect* if one edge is marked as paternal, and the other as maternal, and *unbinned* otherwise. Vertex *v* is then classified as *correct* if all of its resolved links are either true or unbinned, *unbinned* if all resolved links are unbinned, or *incorrect* otherwise. Figure 3 shows the number of correct, incorrect, and unbinned vertices for HUMAN for completely- and partially-resolved vertices in the LJA assembly graph.

**Figure 3.**
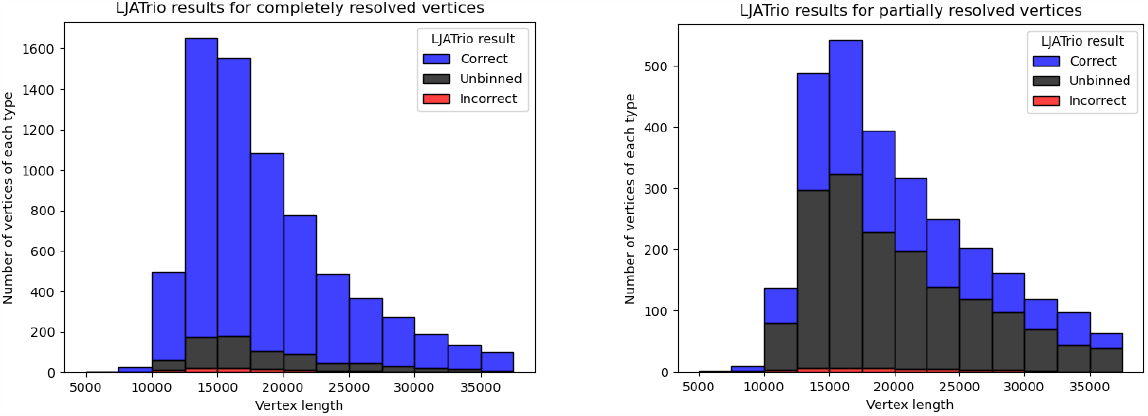
LJATrio validation results. The LJATrio binning results for completely **(left)** and partially **(right)** SpLitter-resolved vertices for the HUMAN dataset. The x-axis shows the vertex length in the LJA assembly graph. The bin width in this histogram is 2500 bp.

#### Coverage effects on the repeat resolution

For the HUMAN dataset, out of 6788 branching vertices that were neither completely nor partially resolved 6241 are uncovered, i.e. none of the in-edge and out-edge pairs share at least *abs_thr* barcodes. In order to analyze, how linked-read coverage affects the outcome of the SpLitteR repeat resolution procedure, we downsampled the larger HUMAN+ dataset (which in total contains ∼5,579 million barcoded TELL-Seq reads) to 10%, 20%, …, 80%, and 90% of all barcodes. As shown in Figure 4, the number of completely resolved vertices rapidly increases, while the number of uncovered vertices decreases with the increase in coverage. However, the rate of this increase slows down after 80% of barcodes are utilized, suggesting that a further increase in coverage is unlikely to significantly improve the assembly quality.

**Figure 4.**
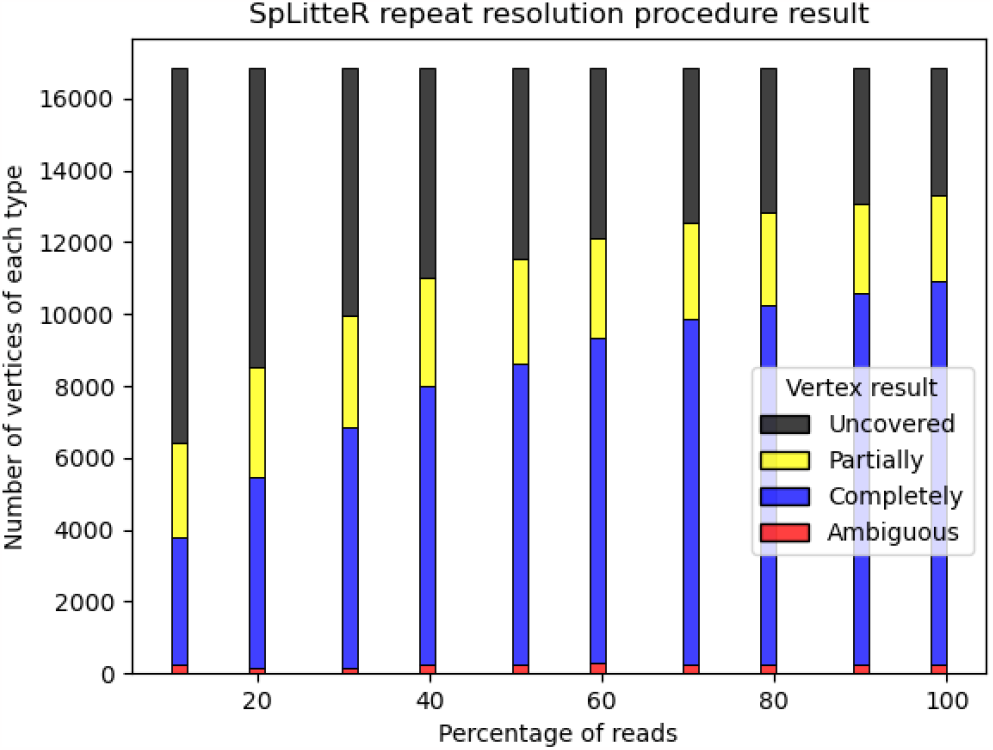
Repeat resolution results for downsampled HUMAN+ dataset. The x-axis shows the selected percentage of HUMAN+ dataset barcodes. Barplots show the number of completely resolved (blue), partially resolved (yellow), uncovered (black), and ambiguous (red) vertices identified by the SpLitteR repeat resolution procedure.

### SHEEP dataset benchmark

Below we describe benchmarking of SpLitteR, ARKS 1.2.4 [15], and SLR-superscaffolder 0.9.1 [17] on the SHEEP dataset. Unlike the HUMAN dataset which was assembled using LJA, the SHEEP dataset was assembled using metaFlye v 2.9-b1768 [9] since LJA was not designed for metagenomic assemblies. In the case of SpLitteR, we benchmarked an assembly formed by sequences of edges in the simplified assembly graph generated by the SpLitteR. In the cases of ARKS and SLR-superscaffolder, we benchmarked their assemblies based on the scaffolds provided by metaFlye.

We used QUAST-LG [19] to compute various reference-free metrics of the resulting assemblies. Table 4 illustrates that all assemblers have similar contiguity with ARKS resulting in the largest NG50, NG25, and auNG metrics for the SHEEP dataset.

**Table 4.** QUAST results for the SHEEP dataset.

| Tool | metaFlye<br>(edges) | metaFlye<br>(scaffolds) | metaFlye<br>+ ARKS | metaFlye +<br>SLR-super<br>scaffolder | metaFlye<br>(edges) +<br>SpLitter |
| --- | --- | --- | --- | --- | --- |
| # contigs ($\geq 0$<br>bp) | 132288 | 104107 | **103321** | 104107 | 130974 |
| # contigs (≥<br>50kbp) | 39661 | 40226 | 39851 | 40266 | 39595 |
| Total length<br>($\geq 50$ kbp) | 6.21 Gbp | 6.31 Gbp | **6.32 Gbp** | 6.31 Gbp | 6.22 Gbp |
| Largest contig | 5,897,638 | 5,953,377 | **5,988,140** | 5,953,377 | 5,897,528 |
| NG50 | 115660 | 119205 | **121777** | 119205 | 116285 |
| NG25 | 388998 | 393824 | **404963** | 393824 | 393824 |
| auNG | 396787 | 399346 | **413319** | 399346 | 401817 |

For the SHEEP dataset, SLR-superscaffolder and SpLitteR scaffolding did not result in any increase in contiguity compared to the initial metaFlye assembly, while ARKS result in a minor increase. Since ARKS and SLR-superscaffolder have very high RAM requirements, we only report SpLitteR results on the high-coverage HUMAN+ dataset.

The suboptimal results demonstrated by SpLitteR assembly compared to ARKS and the base metaFlye assembly stem from the negligible amount of information provided by the assembly graph structure. The metaFlye assembly graph of the SHEEP dataset contains only 923 branching vertices, of which 194 were completely resolved and 433 were partially resolved by SpLitteR. Despite the relative effectiveness of the SpLitteR repeat resolution procedure (more than half the branching vertices were at least partially resolved), graph-agnostic scaffolding performed by ARKS yields more contiguous assembly. The results for the SHEEP dataset demonstrate that the effectiveness of SpLitteR’s scaffolding highly depends on the choice of the baseline assembly graph.

## Discussion

Since linked-read scaffolders, such as SLR-superscaffolder and ARKS, do not utilize the assembly graph information, they have limited applicability to long-read assemblies due to the increased unresolved repeat length compared to short read assemblies. In addition, SLR-superscaffolder utilizes input .bam file to assign barcodes to contigs, while ARKS uses non-unique kmers for the same purpose. In the case of highly repetitive assembly graphs, e.g. constructed from diploid genomes or strain-rich metagenomes, resulting barcode assignments turn out to be inaccurate. Producing inaccurate barcode-edge assignments and ignoring connections between assembly graph edges makes it difficult for both ARKS and SLR-superscaffolder to improve upon baseline LJA assembly in the HUMAN dataset. On the SHEEP dataset, the baseline assembly graph has higher connectivity and thus provides less information that can be used for scaffolding, while the assembly is less repetitive. As a result, ARKS is able to outperform baseline assembly and other scaffolders. Both ARKS and SLR-superscaffolder output scaffolds instead of an assembly graph, which makes it harder to use TELL-Seq in combination with other supplementary sequencing technologies.

SpLitteR uses the assembly graph and employs unique *k*-mer mapping to overcome these shortcomings. For high quality assembly graphs, even the simple SpLitteR repeat resolution algorithm resolves 94.6% repeats in the HUMAN dataset that are bridged by at least two TELL-Seq fragments. However, for the SHEEP metagenome assembly graph with less clear repeat structure, SpLitteR results in less contiguous assembly than ARKS. While SpLitteR is in theory able to take as an input assembly graphs other than metaFlye and LJA, other assemblers which support GFA format are not yet supported. It should also be noted that for the highly repetitive assemblies consisting of long exact repeats with relatively few SNPs, TELL-Seq short read coverage should be quadratic with respect to ultralong read coverage, since two reads in the same barcode should cover two SNP positions in order to provide information, while a single ultralong read is able to cover consecutive SNPs. Despite this limited coverage scalability compared to ultralong reads, using TELL-Seq human genome dataset with 25x coverage was enough to resolve 62% of repeats unresolved by HiFi assembly.

### Conclusions

We developed the SpLitteR tool for scaffolding and haplotype phasing in assembly graphs using linked-reads. Our benchmarking demonstrated that it significantly increases the assembly contiguity compared to the previously developed HiFi assemblers and linked-read scaffolders. We thus argue that linked-reads have the potential to become an inexpensive supplementary technology for generating more contiguous assemblies of large genomes from the initial HiFi assemblies, in line with ONT and Hi-C reads, which were used by the T2T consortium to assemble the first complete human genomes [12, 13]. Since the assembly graph simplification procedure in SpLitteR yields longer contigs as compared to the initial HiFi-based assembly, SpLitteR can be integrated as a preprocessing step in the assembly pipeline with other tools that employ supplementary sequencing technologies, such as Hi-C [10] and Strand-seq [21].

## Materials and Methods

SpLitteR is a tool for resolving repeats in the assembly graph using Tell-Seq data. We assume that the genome defines an (unknown) *genomic traversal* of the assembly graph. Given an incoming edge *e* into a vertex *v*, we define a *follow-up edge next*(*e*) as the edge that immediately follows *e* in this traversal. A vertex in a graph is classified as *branching* if both its in-degree and out-degree exceed 1 (each branching vertex in the graph represents a genomic repeat).

Figure 5 illustrates the SpLitteR workflow. First, SpLitteR maps the barcoded TELL-Seq reads to the edges of the assembly graph, identifies the uniquely mapped reads, and stores their barcodes for each edge (see Section Aligning barcoded reads for details). Given an incoming edge *e* into a branching vertex *v*, SpLitteR attempts to find a follow-up outgoing edge *next*(*e*) by analyzing all linked reads that map to both the in-edge *e* and all out-edges from *v* (see Section Repeat resolution). A vertex is classified as *resolved* if SpLitteR finds a follow-up edge for each incoming edge into this vertex. SpLitteR further simplifies the assembly graph by *splitting* the resolved vertices in such a way that each matched pair of an in-edge and an out-edge is merged into a single edge. Finally, it outputs the results of the repeat resolution procedure both as the set of scaffolds and as the simplified assembly graph. The repeat resolution procedure has both diploid and metagenomic modes.

**Figure 5.**
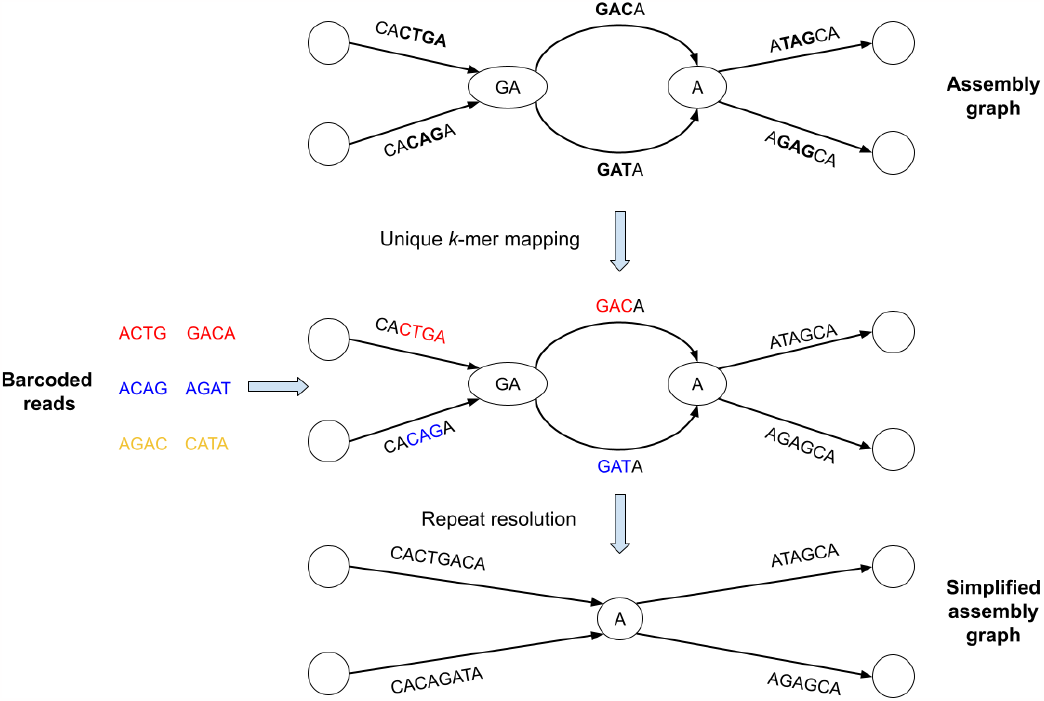
Brief summary of the SpLitteR workflow. In this toy example, the assembly graph is represented as the *multiplex de Bruijn graph* [11] where vertices are labeled by *k*-mers of varying sizes. Reads with the same barcode are represented by the same color (each barcode contains only two reads) Reads are mapped to the assembly graph based on their *unique k*-mers, i.e., *k*-mers which occur only once in the edges of the assembly graph (*k*=3 for this toy example). Since yellow reads do not contain unique 3-mers, they remain unmapped. SpLitteR resolves vertices in the multiplex de Bruijn graph by assigning in-edges to their follow-up out-edges based on the barcode information.

### Data preparation

To generate the TELL-Seq dataset from the HG002 genome, 5ng high molecular weight genomic DNA, extracted from GM24385 (HG002) cells, was used to construct TELL-Seq WGS libraries based on the manufacturer’s user guide for large genome library preparation. These libraries were sequenced as 2×150 paired-end reads on a NovaSeq instrument.

A fecal sample of the SHEEP dataset was taken from a young (<1 year old) wether lamb of the Katahdin breed. DNA was extracted in small batches from approximately 0.5 g per batch using the QIAamp PowerFecal DNA Kit, as suggested by the manufacturer (Qiagen), with moderate bead beating. 5ng DNA was used to construct TELL-Seq WGS libraries based on the manufacturer’s user guide for large genome library preparation (Universal Sequencing Technology). These libraries were sequenced as 2×150 paired end reads on a NovaSeq instrument.

Since homopolymer-compressed HiFi reads have significantly lower error rates than raw HiFi reads, assembly graphs produced by some of the existing HiFi-based assemblers, such as LJA [11], are homopolymer-compressed. Thus, to generate read-to-graph alignments to such assembly graphs, each homopolymer run X…X in each TELL-Seq read was collapsed into a single nucleotide X. Additionally, long dimer repeats (of length 16 and more) were compressed as described in [11]. Assembly produced by the metaFlye was not homopolymer-compressed. Adapters were removed from TELL-Seq barcoded reads using cutadapt v 4.1 [22].

### Representation of assembly graphs

SpLitteR was originally designed to operate on the *multiplex de Bruijn graphs* (*mDBG*) generated by the LJA assembler [11]. However, it also supports arbitrary assembly graphs in the GFA format such as those generated by Verkko [12], Flye / metaFlye [9, 23], Shasta [8] and other genome assembly tools. Since the representation of such graphs in the standard GFA format differs from the mDBG encoding, they require an additional transformation into *DBG-like* graphs (GFA segments correspond to edges and unresolved repeats correspond to vertices).

Let *G* be an arbitrary assembly graph in the GFA format consisting of a set of segments *E*(*G*) and links *L*(*G*). We expand *E*(*G*) into a set *E’*(*G*) of segments *e* from *E*(*G*) and their reverse complements *rc*(*e*). Afterward, we transform *G* into a directed *raw DBG-like* graph *RDG*, with edges *E*(*RDG*) *= E’*(*G*). For every edge *e* in *E’*(*G*), we form vertices *start*(*e*) and *end*(*e*). We then merge those vertices according to the GFA links. For every GFA link (*e*_*1*_, *e*_*2*_) from *L*(*G*), we construct *link edges* between *end*(*e*_*1*_) and *start*(*e*_*2*_), and between *end*(r*c*(*e*_*2*_)) and *start*(*rc*(*e*_*1*_)).

We say that *contraction* of an edge (*v, w*) is the merging of *v* and *w* into a single vertex *u*, followed by the removal of the loop-edge resulting from this merging. Then DBG-like graph DG is obtained by contracting every link edge in the raw DBG-like graph *RDG*. For every vertex *v* in *RDG* we store GFA links *L*(G, *v*) which were contracted into *v* in order to retain the connectivity information from *G*.

For metaFlye graphs, we contract edges that were classified as repetitive by metaFlye. In the resulting *contracted* assembly graph *CG* every non-leaf vertex represents an unresolved and possibly inexact repeat from the input assembly.

### Aligning barcoded reads

SpLitteR maps short reads from a linked-read library to the edges of the contracted assembly graph using the *k*-mer-based alignment approach originally developed for mapping Hi-C reads [24]. First, we collect unique (occurring only once) *k*-mers (default value *k*=31) in the edge sequences of the contracted assembly graph and record their edge positions in this graph. In the metagenomic mode, we map a barcoded read-pair to an edge if both reads from the pair contain at least one unique *k*-mer from this edge. In the diploid mode, we relax the mapping requirement to a single *k*-mer from the entire read-pair, as most *k*-mers in the phased HiFi assembly graph are repetitive. Our analysis has shown that most 31*-*mers in assembly graphs constructed for a single human haplome or for a metagenome are unique, which ensures a sufficient number of unique *k*-mers in heterozygous regions of the assembly graph. For every edge *e* in the contracted graph, we store the barcode-set *barcodes*(*e*), comprising barcodes of all reads mapped to *e*.

### Repeat resolution

In order to resolve a potential repeat, for each incoming edge *e* in vertex *v*, we aim to find the edge *next*(*e*) in the contracted assembly graph. Our base assumption is that consecutive edges *e* and *next*(*e*) in the genomic traversal have similar (overlapping) barcode-sets, as many fragments that contain a suffix of *e* also contain a prefix of *next*(*e*).

Given an in-edge *e* into a vertex *v* and an out-edge *e’* from this vertex, we define *overlap*(*e, e’*) as the size of the intersection of barcode-sets *barcodes*(*e*) and *barcodes*(*e’*). For an incoming edge *e* into a branching vertex *v*, we define *overlap*_*1*_(*e*) (*overlap*_*2*_(*e*), respectively) as the largest (second largest) among all values of *overlap*(*e,e’*) among all outgoing *e’* edges from *v*. We refer to an out-edge *e’* with the largest value of *overlap*(*e,e’*) as *e\** (ties are broken arbitrarily).

We use a simple decision rule for finding a candidate edge *next*(*e*) for each in-edge *e*. Specifically, we select the edge *e\** if *overlap*_*1*_(*e*) ≥ *rel_thr * overlap*_*2*_(*e*) and *overlap*_*1*_(*e*) ≥ *abs_thr* (default values *rel_thr =* 2 and *abs_thr* = 2). If these conditions hold, we say that the edge-pair *(e, e\**) is a *candidate link* for vertex *v*. If for every edge *e* incident to *v*, |*barcodes*(*e*)| < *abs_thr*, we say that *vertex* is *uncovered*.

### Assembly graph simplification and scaffolding

We say that a vertex *v* is *partially resolved* if candidate links constitute a non-empty *matching M* between in-edges and out-edges of *v*, and *completely resolved* if this matching is a *perfect matching*. If *v* is not completely resolved, partially resolved, or uncovered, we say that *v* is *ambiguous*.

For LJA assembly graphs, we perform a splitting procedure for every completely or partially resolved vertex *v* in order to simplify the assembly graph for possible later scaffolding. For every completely or partially resolved vertex, the set of candidate links (*e, next*(*e*)) comprises a matching. For every candidate link (*e, next(e*)) in this matching we create a new vertex *v*_*e*_, such that *e* is the only in-edge into *v*_*e*_, and *next(e)* is the only out-edge out of *v*_*e*_. If *v* is completely resolved, we then remove it from the graph. After performing the splitting procedure for every completely or partially resolved vertex, we condense the non-branching paths *Paths* in the contracted graph *CG*. The resulting graph is outputted in the GFA format.

For all other assembly graphs, the non-branching paths from *Paths* are outputted as scaffolds without changing the original assembly graph *G*. For every pair of scaffolded edges in *CG* (*e, next*(*e*)), if there is a unique path in *G* from *end*(*e*) to *start*(*next*(*e*)), the sequence of this path is inserted between *e* and *next*(*e*) in the resulting scaffold. Otherwise, a sequence of *N* characters of length *Distance*_*G*_ (*e*_*1*_, *e*_*2*_) is inserted between *e* and *next*(*e*), where *Distance*_*G*_ (*e, next*(*e*)) is the distance in *G* from *end*(*e*) to *start*(*next*(*e*)).

## Declarations

### Ethics approval and consent to participate

Not applicable

## Consent for publication

Not applicable

## Availability of data and materials

SpLitteR is implemented in C++ as a part of the freely available SPAdes package and is available at https://github.com/ablab/spades/releases/tag/splitter-preprint

The sequencing reads for the HUMAN dataset are available in the NCBI BioProject database under accession number SRX7264481. The remaining reads for the HUMAN+ and SHEEP datasets generated in this study have been submitted to the NCBI BioProject database under accession number PRJNA956112. Baseline LJA assembly and trio binning results for the HUMAN+ dataset are available at https://figshare.com/articles/dataset/HG002/21678842. Baseline metaFlye assembly for the SHEEP dataset is available at https://figshare.com/articles/dataset/SHEEP/22864043.

## Authors’ contributions

Conceptualization, IT and AK; methodology, IT, PP and AK.; software, IT and AK.; writing—original draft preparation, IT; writing—review and editing, IT, ZC, PP and AK.; data acquisition, ZC; supervision, AK.; funding acquisition, AK. All authors have read and agreed to the published version of the manuscript.

## Acknowledgements

The research was carried out in part by computational resources provided by the Resource Center “Computer Center of SPbU.”

## Tables

**Table 1. Information about TELL-Seq datasets.** Mean fragment length and mean number of fragments per container statistics were evaluated based on the T2T reference for the HUMAN dataset, and the metaFlye assembly for the SHEEP dataset. A fragment is defined as the set of 2 or more paired-end reads with the same barcode aligned to the same long (>500kbp) scaffold. Mean fragment length is the mean distance from the start of the leftmost aligned read in a fragment to the rightmost aligned read in a fragment.

**Table 2. QUAST results for the HUMAN dataset.**

**Table 3. Repeat resolution statistics for the HUMAN, HUMAN+, and SHEEP datasets.**

**Table 4. QUAST results for the SHEEP dataset.**

## References

1. McElwain MA, Zhang RY, Drmanac R, Peters BA. Long Fragment Read (LFR) Technology: Cost-Effective, High-Quality Genome-Wide Molecular Haplotyping. Methods Mol Biol. 2017;1551:191–205.

2. Chen Z, Pham L, Wu T-C, Mo G, Xia Y, Chang PL, et al. Ultralow-input single-tube linked-read library method enables short-read second-generation sequencing systems to routinely generate highly accurate and economical long-range sequencing information. Genome Res. 2020;30:898–909.

3. Callahan BJ, Grinevich D, Thakur S, Balamotis MA, Yehezkel TB. Ultra-accurate microbial amplicon sequencing with synthetic long reads. Microbiome. 2021;9:130.

4. Bishara A, Moss EL, Kolmogorov M, Parada AE, Weng Z, Sidow A, et al. High-quality genome sequences of uncultured microbes by assembly of read clouds. Nat Biotechnol. 2018;36:1067–75.

5. Tolstoganov I, Bankevich A, Chen Z, Pevzner PA. cloudSPAdes: assembly of synthetic long reads using de Bruijn graphs. Bioinformatics. 2019;35:i61–70.

6. Weisenfeld NI, Kumar V, Shah P, Church DM, Jaffe DB. Direct determination of diploid genome sequences. Genome Res. 2017;27:757–67.

7. Nurk S, Walenz BP, Rhie A, Vollger MR, Logsdon GA, Grothe R, et al. HiCanu: accurate assembly of segmental duplications, satellites, and allelic variants from high-fidelity long reads. Genome Res. 2020;30:1291–305.

8. Shafin K, Pesout T, Lorig-Roach R, Haukness M, Olsen HE, Bosworth C, et al. Nanopore sequencing and the Shasta toolkit enable efficient de novo assembly of eleven human genomes. Nat Biotechnol. 2020;38:1044–53.

9. Kolmogorov M, Bickhart DM, Behsaz B, Gurevich A, Rayko M, Shin SB, et al. metaFlye: scalable long-read metagenome assembly using repeat graphs. Nat Methods. 2020;17:1103–10.

10. Cheng H, Concepcion GT, Feng X, Zhang H, Li H. Haplotype-resolved de novo assembly using phased assembly graphs with hifiasm. Nat Methods. 2021;18:170–5.

11. Bankevich A, Bzikadze AV, Kolmogorov M, Antipov D, Pevzner PA. Multiplex de Bruijn graphs enable genome assembly from long, high-fidelity reads. Nat Biotechnol. 2022;40:1075–81.

12. Rautiainen M, Nurk S, Walenz BP, Logsdon GA, Porubsky D, Rhie A, et al. Telomere-to-telomere assembly of diploid chromosomes with Verkko. Nat Biotechnol. 2023. 10.1038/s41587-023-01662-6.

13. Nurk S, Koren S, Rhie A, Rautiainen M, Bzikadze AV, Mikheenko A, et al. The complete sequence of a human genome. Science. 2022;376:44–53.

14. Kuleshov V, Snyder MP, Batzoglou S. Genome assembly from synthetic long read clouds. Bioinformatics. 2016;32:i216–24.

15. Coombe L, Zhang J, Vandervalk BP, Chu J, Jackman SD, Birol I, et al. ARKS: chromosome-scale scaffolding of human genome drafts with linked read kmers. BMC Bioinformatics. 2018;19:234.

16. Afshinfard A, Jackman SD, Wong J, Coombe L, Chu J, Nikolic V, et al. Physlr: Next-Generation Physical Maps. DNA. 2022;2:116–30.

17. Guo L, Xu M, Wang W, Gu S, Zhao X, Chen F, et al. SLR-superscaffolder: a de novo scaffolding tool for synthetic long reads using a top-to-bottom scheme. BMC Bioinformatics. 2021;22:158.

18. Wenger AM, Peluso P, Rowell WJ, Chang P-C, Hall RJ, Concepcion GT, et al. Accurate circular consensus long-read sequencing improves variant detection and assembly of a human genome. Nat Biotechnol. 2019;37:1155–62.

19. Mikheenko A, Prjibelski A, Saveliev V, Antipov D, Gurevich A. Versatile genome assembly evaluation with QUAST-LG. Bioinformatics. 2018;34:i142–50.

20. Antipov D, Bankevich A, Bzikadze A. LJATrio development branch. GitHub. 2022. https://github.com/AntonBankevich/LJA/tree/LJAtrio. xAccessed 31 Oct 2022.

21. Porubsky D, Ebert P, Audano PA, Vollger MR, Harvey WT, Marijon P, et al. Fully phased human genome assembly without parental data using single-cell strand sequencing and long reads. Nat Biotechnol. 2021;39:302–8.

22. Martin M. Cutadapt removes adapter sequences from high-throughput sequencing reads. EMBnet.journal. 2011;17:10.

23. Kolmogorov M, Yuan J, Lin Y, Pevzner PA. Assembly of long, error-prone reads using repeat graphs. Nat Biotechnol. 2019;37:540–6.

24. Cheng H, Jarvis ED, Fedrigo O, Koepfli K-P, Urban L, Gemmell NJ, et al. Haplotype-resolved assembly of diploid genomes without parental data. Nat Biotechnol. 2022;40:1332–5.

